# Calcium-dependent reactive oxygen formation and blood-brain barrier breakdown by NOX5 limits post-reperfusion outcome in stroke

**DOI:** 10.1101/359893

**Authors:** Ana I Casas, Pamela WM Kleikers, Eva Geuss, Friederike Langhauser, Javier Egea, Manuela G Lopez, Christoph Kleinschnitz, Harald HHW Schmidt

**Author notes:** These authors contributed equally to this word (Ana I Casas, Pamela WM Kleikers). These authors are joint senior authors on this work (Christoph Kleinschnitz, Harald HHW Schmidt).

## Abstract

Ischemic stroke is a predominant cause of disability worldwide, with thrombolytic or mechanical removal of the occlusion being the only therapeutic options. Reperfusion bears the risk of an acute deleterious calcium-dependent breakdown of the blood-brain-barrier. Its mechanism, however, is unknown. Here we identify type 5 NADPH oxidase (NOX5), a calcium-activated, reactive oxygen species (ROS)-forming enzyme as missing link. Using a humanised knock-in mouse model and *in vitro* in organotypic cultures, we find re-oxygenation or calcium overload to increase brain ROS levels in a NOX5-dependent manner. *In vivo*, post-ischemic ROS formation, infarct volume and functional outcomes were worsened in NOX5 knock-in mice. Of clinical and therapeutic relevance, in a human blood-barrier model pharmacological NOX inhibition also prevented acute re-oxygenation induced leakage. Our data therefore identify NOX5 as sufficient to induce acute post-reperfusion calcium-dependent blood-brain-barrier breakdown. We suggest urgent clinical validation by conducting protective post-stroke re-canalisation in the presence of a NOX inhibitor.

## Introduction

Ischemic stroke represents one of the most frequent causes of death and leading cause of disability worldwide [1]. In the absence of any effective neuroprotective principle, thrombolytic or mechanical removal of the occlusion remains the only current therapeutic option [2]. In 11% of the mechanically re-canalysed patients, however, serious complications occur including new emboli formation, vasospasm, intracranial haemorrhage and stent dislocation or occlusion [3]. Similarly, thrombolysis using tissue plasminogen activator (rt-PA) correlates with an early opening of the blood-brain barrier (BBB)^4^, which may lead to pathologic processes such as edema and hemorrhage transformation^4^ together with excessive reactive oxygen species (ROS) production, neuronal death and therefore worse patient prognosis [4]. Moreover, an increase in intracellular calcium represents one of the earliest events during an ischemic stroke and is considered to trigger many of the downstream processes which promote blood-brain barrier leakage and brain oedema, the leading cause of death after an ischemic stroke [2,5].

With respect to ROS, NADPH oxidases are considered a primary and quantitatively relevant source in different ischemic conditions [5]. Of particular interest is NADPH oxidase type 5 (NOX5), the calcium-activated isoform of this enzyme family and widely expressed in the endothelium [6], testis and white blood cells [7]. We therefore hypothesised that NOX5 may be the missing mechanistic link between post-reperfusion calcium overload and early BBB opening. To validate this target, we examined both a pharmacological approach, using a panel of the most advanced inhibitor compounds in a human *in vitro* BBB model, and a genetic approach, using the most common *in vivo* stroke model in mice and following the STAIRS quality criteria. Since NOX5 is missing from the mouse genome, we also generated a novel humanised mouse model expressing the human NOX5 gene in the neutral hypoxanthine phospho-ribosyl-transferase *(Hprt)* locus.

## Results

### Generation and validation of the humanised NOX5 KI mouse model

In humans, *Nox5* is broadly expressed in blood vessels, primarily smooth muscle cells, microvascular endothelial cells, fibroblasts [6,8,9] but also in testis [8] and spleen, specifically in monocytes, macrophages [7,10], B- and T-cells [11]. To mimic the physiological human expression pattern of NOX5 in mice, we created a mouse line called NOX5 KI (knock-in) bearing the human *NOX5* gene in the *Hprt* locus and under control of the *Tie2* promoter (Fig 1A-B). The *Tie2* promoter physiologically regulates endothelial and hematopoietic gene expression in human cells [12,13]. Therefore, using this promoter we expected our transgenic mouse model to mimic physiological human *NOX5* expression, both in endothelial and white blood cells.

**Fig 1.**
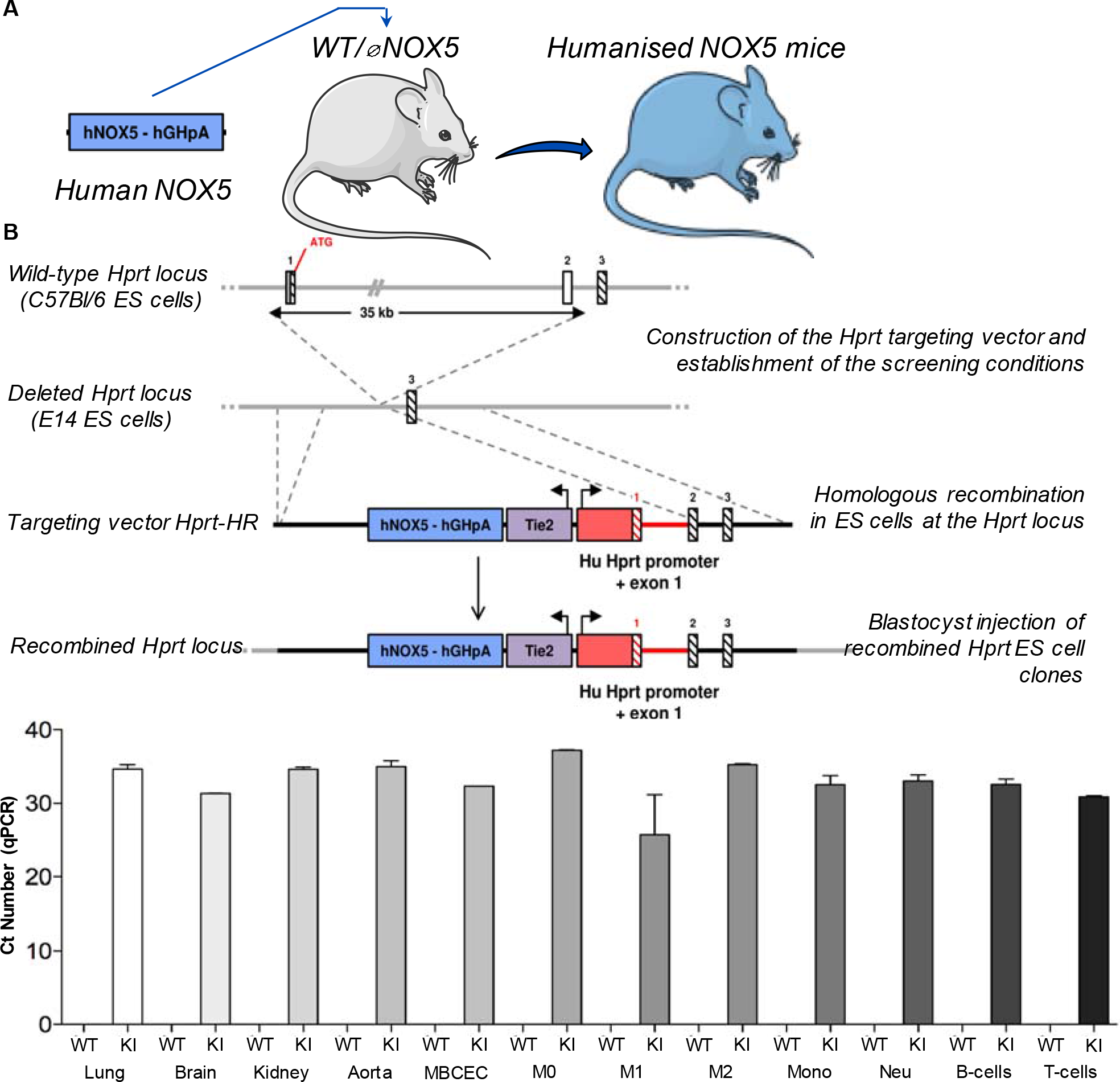
Generation and validation of the NOX5 KI mouse. (A) Representative scheme of the humanised NOX5 mice. *NOX5* gene is now located on the X chromosome resulting in females with a double copy of the gene (NOX5^KI/KI^) while gene load will be half in male mice (NOX5^KI/Y^) (B) Schematic representation of the construction of the humanised NOX5 knock-in (KI) mice. First, the wild type and deleted Hprt constructs are shown. Third, targeting vector to insert NOX5: human NOX5 cDNA (blue box) is coupled to the human Hprt promoter (red box) and exons 1 and 2. The human Hprt promoter is under control of the Tie2 gene (purple box) to create NOX5 expression. Hatched black and red boxes represent murine and human *Hprt* exons, respectively. Solid line represents intronic sequences. Diagram is not depicted to scale. (C) *NOX5* gene expression was measured by qPCR (Ct numbers) in different organs: lung, brain, kidney, aorta and mice brain capillary endothelial cells (MBCEC); macrophages from bone marrow: undifferentiated (M0), inflammatory (M1) and active (M2); and hematopoietic cells isolated from spleen: monocytes (Mono), neutrophils (Neu), B-cells and T-cells. Tissues from WT mice did not show *NOX5* gene expression, while significant expression was detected in NOX5 KI mice. CT values of n = 4 are shown.

We examined *NOX5* gene expression in different organs and cells from the offspring mice. Tissues from WT mice did not show any detectable *NOX5* gene expression, whilst significant expression was detected in all samples from KI mice (Fig 1C). Similarly, genotyping agarose electrophoresis showed the presence of *NOX5* cDNA in lung, kidney, brain, aorta and white blood cells (monocytes, lymphocytes, T and B-cells; Fig S1). These data suggested that, with the exception of testis, our NOX5 KI mice can be considered a humanised mouse mode showing no specific overall phenotype in comparison with non-transgenic wild type mice in terms of lifespan, cognition or neuro-motor function.

### *In vitro*, NOX5 causes acute, calcium-dependent post-reoxygenation ROS formation

Distinct NOX isoforms, i.e. NOX2 and NOX4, have been described as key players in stroke ROS-dependent patho-mechanism [5,14], however, no specific role of NOX5 has been defined so far. Thus, to examine whether NOX5 causes acute cerebral post-reperfusion/re-oxygenation ROS formation, we first studied *in vitro* organotypic hippocampal cultures (OHC) of NOX5 KI mice. When we subjected OHCs to oxygen and glucose deprivation for 15 min (OGD; Fig 2A) and assessed kinetics of ROS formation at 0, 15, 30 min and 1, 2 and 4 h post-OGD, we detected an acute NOX5-dependent ROS production within the first half hour that was not present in WT mice (Fig 2B-C). At later time-points, ROS were unchanged presumably due to the NOX4 isoform [15]. To confirm that this observation was indeed due to calcium-induced activation of NOX5, we re-examined ROS generation in OHCs from NOX5 KI and WT mice in the absence or presence of the calcium ionophore A23187 (10μM), but without OGD. A23187 is a mobile ion-carrier which forms stable complexes with divalent cations, mainly Mn^2+^ and Ca^2+^ leading to increase in intracellular levels of Ca^2+^ [16]. Indeed, we observed the same acute ROS surge as after OGD and in a strictly NOX5-dependent manner (Fig. 2D-E). Thus, these data suggest that NOX5-dependant ROS formation is involved in early stages of post-ischemia damage via Ca^2+^ overload.

**Fig 2.**
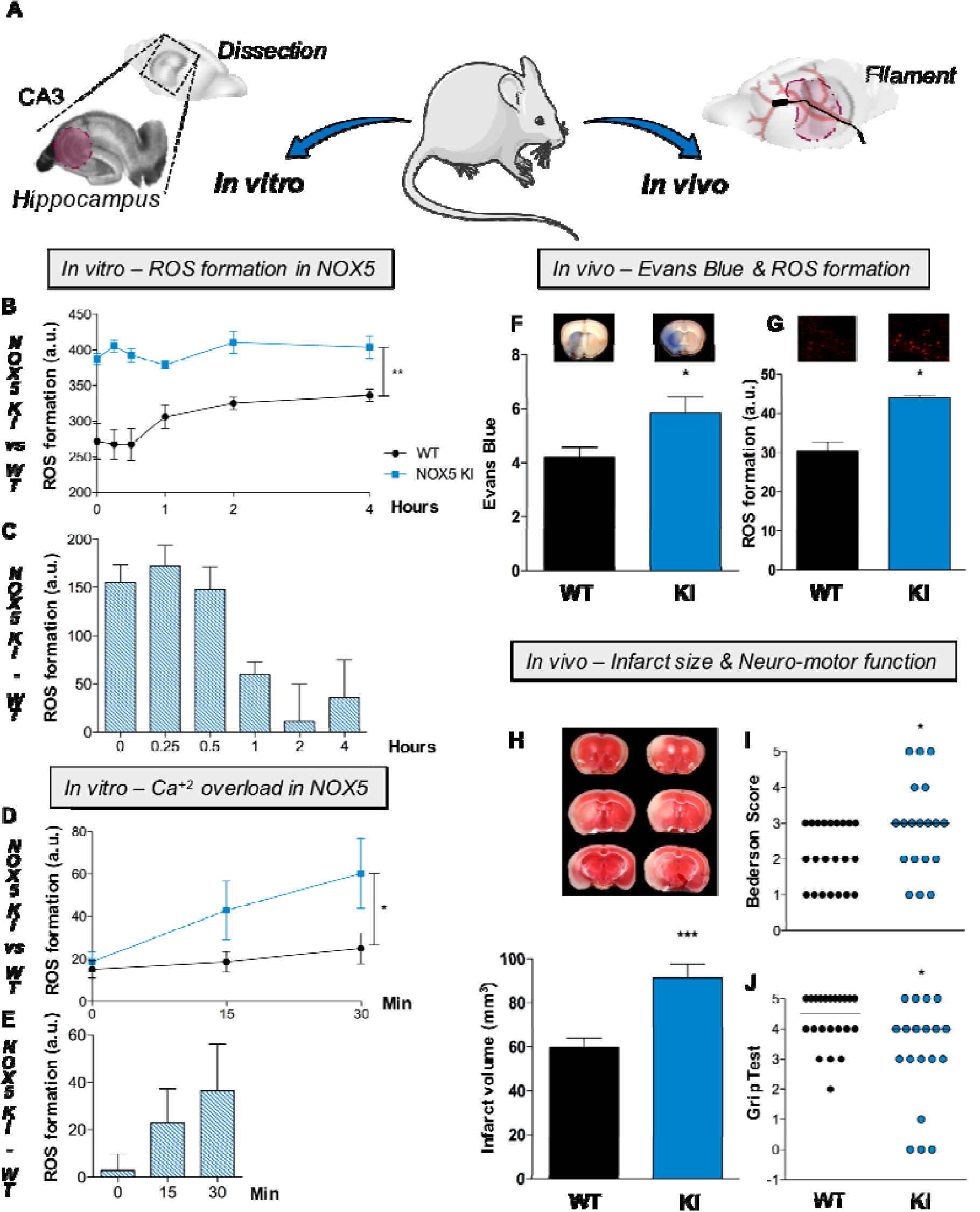
Increased *in vitro* ROS production and Ca^2+^ overload in NOX5 KI mice lead to enlarged stroke size, blood-brain barrier damage and aggravate neuro-motor function. (A) Two different brain ischemia models were used. *In vitro*, mice organotypic hippocampal cultures (OHCs) were subjected to oxygen and glucose deprivation (OGD) followed by the re-oxygenation (Re-Ox) period. *In vivo*, occlusion of the middle cerebral artery was performed in WT and NOX5 KI mice. (B) ROS production measured after OGD in organotypic hippocampal cultures was higher in NOX5 KI mice (blue, n = 6) versus NOX5 WT mice (black, n = 6) starting directly from the OGD period until 4 h post-OGD (** P < 0.01). (C) Difference in ROS production between NOX5 KI (blue, n = 6) and WT (black, n = 6) mice showed a NOX5-dependant effect within the first hour post-OGD (0, 0.25, 0.5 and 1h). (D) Addition of the ionophore A23187 (10µM) showed a Ca^2+^-dependent increased of ROS formation in NOX5 KI mice (blue, n = 3) while no changes were detected in WT mice (black, n = 3) (* P < 0.05). (E) Difference in ROS production between NOX5 KI (blue, n = 3) and WT (black, n = 3) mice showed a direct link between Ca^+2^ overload and NOX5 induction at 15 and 30 min post-A23187. (F) Evans blue staining 24h post-tMCAO showed blood-brain barrier damage in the contralateral brain side in both WT (black, n = 21) and NOX5 KI (blue, n = 21) mice (* P < 0.05). Representative pictures are shown above the graphs. (G) ROS production as measured by DHE staining 24h after stroke induction was increased in NOX5 KI (blue, n = 4) versus NOX5 WT (black, n = 4) mice (** P < 0.01). Representative pictures are shown above the graphs. (H) Infarct size as measured by TTC staining showed increased infarct sizes in NOX5 KI (blue, n = 16) versus NOX5 WT mice (black, n = 21) (*** P < 0.001). Neurological behaviour testing showed worse neurological functioning in NOX5 KI (blue circles, n = 19) versus WT (black circles, n = 22) mice with worse (I) Bederson and (J) Grip test scores (* P < 0.05).

### *In vivo*, ROS formation and blood-brain barrier breakdown are NOX5 dependent and worsen neurological outcome

Having established the role of NOX5 after re-oxygenation leading to calcium-dependent ROS formation *in vitro*, we wanted to examine the role of NOX5 in a stroke *in vivo* animal model. We performed 1h of transient occlusion of the middle cerebral artery (tMCAO) on 8-16 weeks-old mice, followed by 24h of reperfusion (Fig 2A) for later histological and neurological assessment.

Reperfusion injury, comparable to *in vitro* re-oxygenation damage, leads to abnormally permeable capillary bed resulting from a disruption of the BBB [4]. Since NOX5 seems to be part of this subsequent post-stroke event, we assessed the role of NOX5 in BBB disruption by quantifying post-stroke edema formation using the extravasation of the macromolecular vascular tracer, Evans blue, into the brain parenchyma. BBB leakage was significantly increased in brains from NOX5 KI mice when compared to their WT littermates, indicating a key role of NOX5 in post-stroke BBB integrity (Fig 2F). Second, to examine the mechanistic link between this observation to NOX5 activity, we measured ROS formation using dihydroethidium (DHE)-stained cryosections of NOX5 KI and WT mice. Again, ROS generation was significantly increased in the infarcted brains of NOX5 KI mice compared to their controls (Fig 2G). Thus, post-stroke NOX5-dependent impairment of the BBB is related to enhanced *in vivo* ROS formation.

To further evaluate whether NOX5-dependent ROS formation and impairment of the BBB were linked to a worsened outcome, we assessed 24h post-stroke infarct sizes. Again, TTC-stained brain sections showed a significant increase of infarct volume in NOX5 KI compared to WT mice (Fig 2H). As a clinical translational parameter, we evaluated neuro-motor function via the Bederson score and Grip tests, two key outcome parameters. We saw that both scores showed worsened neuro-motor function in NOX5 KI animals compared to WT (Fig 2G-I). Thus, acute post-re-oxygenation Ca^2+^-dependent ROS formation, BBB breakdown, increased infarct size and worsened neuro-motor outcome appear to be mechanistically linked to NOX5.

### Blood pressure does not contribute as a confounder to the NOX5 KI phenotype in stroke

With respect to our *in vivo* data, our NOX5 KI approach may have lead to a systemic constitutively elevated endothelial ROS formation, impaired vasodilatory nitric oxide signaling and increased blood pressure. Hypertension can weaken brain arteries and worsen stroke outcome [16] and thus lead to an overestimation of the *in vivo* impact of NOX5 via acutely impairing the blood-brain barrier post-reperfusion. We therefore also assessed the 24h blood pressure in our mice using telemetric blood pressure devices allowing non-invasive recording. Neither systolic (Fig 3A), diastolic (Fig 3B) nor mean arterial pressure (Fig 3C) were different between WT and NOX5 KI mice suggesting that no systemic hypertensive phenotype contributes to the worsened post-reperfusion *in vivo* outcome in NOX5 KI mice. Similarly, when expressing human NOX5 in a non-physiological smooth muscle cell-specific manner basal blood pressure was also normal [17].

**Fig 3.**
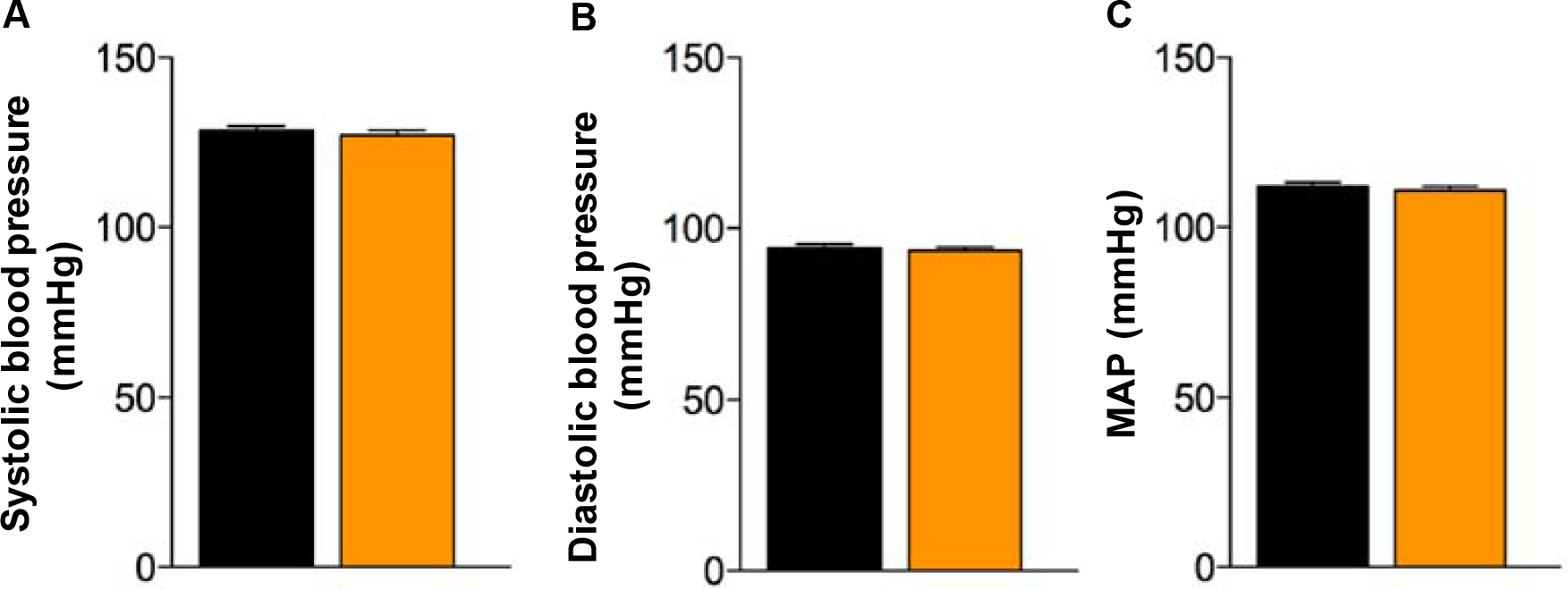
No role of NOX5 in blood pressure. (A) Systolic blood pressure was not different between WT (black, n = 19) and NOX5 KI mice (orange, n = 20). (B) Diastolic blood pressure and (C) mean arterial pressure (MAP) did not differ between WT (black, n = 19) and NOX5 KI (orange, n = 20) mice during day-time (07:00-18:00) or night-time (18:00-06:00).

### The role for NOX5 KI in ischemia reperfusion is specific to the brain

In addition, our NOX5 KI approach may worsen ischemia-reperfusion outcomes in different organs and thus play no brain or BBB-specific role. Indeed, other NADPH oxidases, i.e. NOX 1 and NOX2, have been described as key players in different ischemic diseases such as myocardial ischemia-reperfusion [18], retinopathy [19] or diabetic nephropathy [20,21]. Moreover, Nox5 has been found to be upregulated in human blood vessels after myocardial infarction [22] but its role here remains unclear. We therefore investigated three different *in vivo* mouse models related to highly relevant ischemic disease conditions: hindlimb ischemia (HL, peripheral artery disease), myocardial infarction (MI) and ischemia-reperfusion (IR) of the heart.

To mimic human peripheral artery disease, we used a mouse model of hindlimb ischemia by subjecting 8-16 weeks-old NOX5 KI and WT mice to permanent ligation of the femoral artery. Sprouting of new capillaries (angiogenesis) is commonly used as the major histological outcome post-ligation [23]. Therefore, we measured capillary density using CD31 staining in the gastrocnemius and adductor muscles as the major angiogenesis parameter 4-weeks post-occlusion. CD31 staining was comparable between WT and NOX5 KI mice (Fig 4A-B). We also measured blood flow in both groups and saw a significant drop after surgery, which then gradually increased again within 4-weeks recovery. However, we detected no statistically significant difference in blood flow between the NOX5 KI and WT mice lines (Fig 4C).

**Fig 4.**
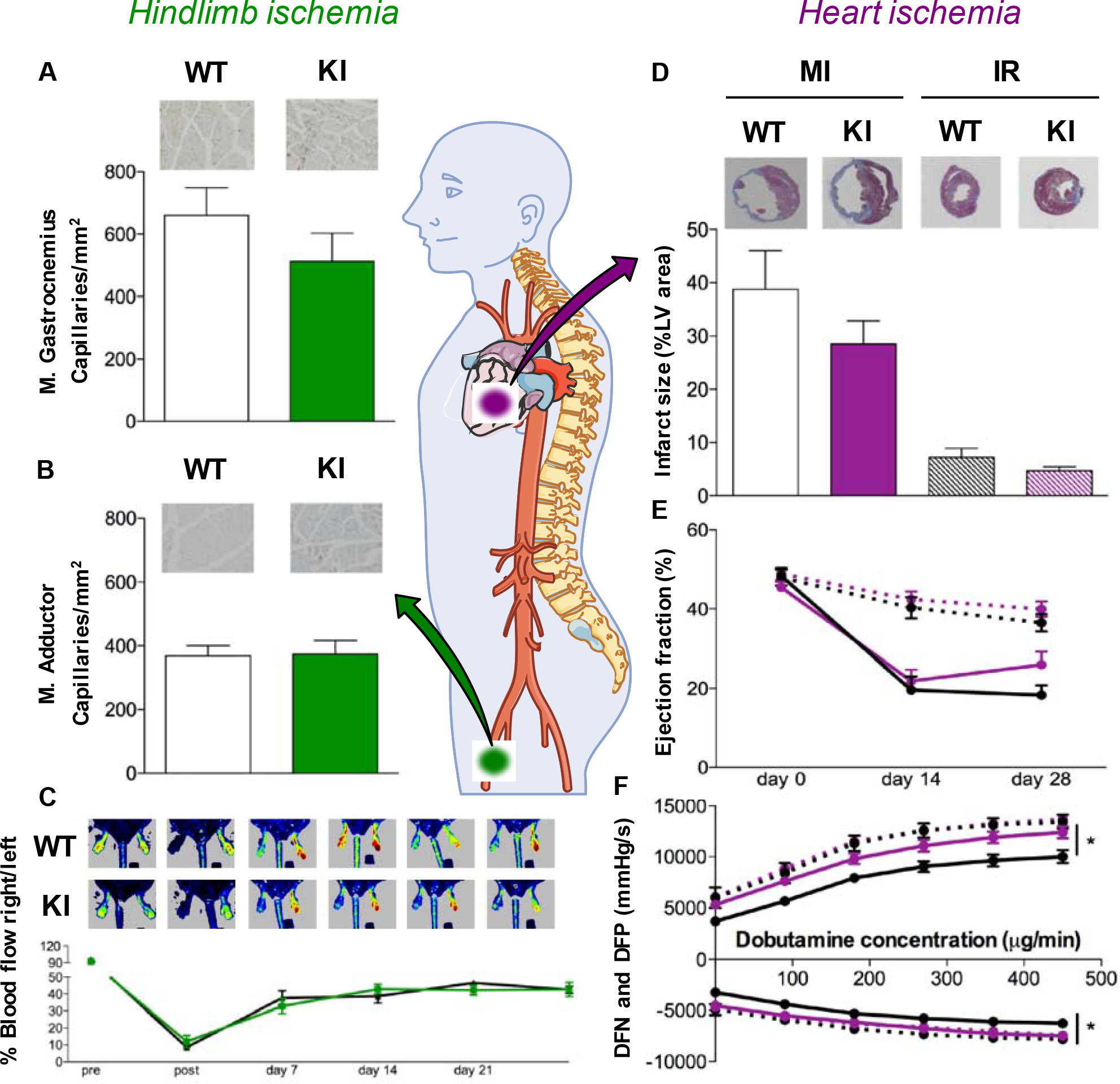
No role of NOX5 in three different cardiovascular ischemic models. Capillary density (capillaries/mm^3^) as measured by CD31 staining showed no difference between NOX5 KI (green) and WT (black) mice 4 weeks after ligation of the femoral artery in (A) gastrocnemius (KI, n = 13; WT, n = 13) (B) and adductor muscles (KI, n = 14; WT, n = 12). (C) Blood flow restoration after ligation of the left femoral artery was not different between WT (black, n = 13) and NOX5 KI (green, n = 10) mice. (D) Infarct size as measured by AZAN staining was not different between WT (black, n = 16) and NOX5 KI mice (purple, n = 15) in the myocardial infarction model. Similarly, no difference was shown in the transient ischemia-reperfusion heart model (hatched black, n = 13; hatched purple, n = 17). (E) Ejection fraction decreased 2 and 4 weeks after permanent (full lines) or transient (dashed lines) heart ischemia. No differences were found between WT (black, n = 16; hatched, n = 12) and NOX5 KI mice (purple, n = 14; hatched, n = 16). (F) Functional measurements of the heart showed better contractile and relaxing properties of the NOX5 KI hearts 4 weeks after permanent ligation (full purple lines, n = 15) compared to WT (full black lines, n = 13) (* P < 0.05). In transient ischemia (dashed lines; WT, n = 12; KI, n = 17), no difference was seen.

To further investigate the role of NOX5 in myocardial infarction and ischemia-reperfusion of the heart, we subjected adult NOX5 KI and WT mice (8-16 weeks old) to a permanent or transient (45 min) occlusion of the left descending coronary artery. Four weeks after a MI or IR, we assessed infarct sizes by AZAN stain, which showed no differences between NOX5 KI and WT mice (Fig 4D). Ejection fraction at 2 and 4 weeks post-MI or post-IR, a common functional parameter, did not differ between NOX5 KI and WT mice (Fig 4E). For an additional clinical parameter, we measured heart contractility in response to increasing concentrations of dobutamine, a sympathomimetic drug, which stimulates heart contraction. NOX5 KI mice subjected to myocardial infarction showed slightly enhanced contractility and relaxation effects compared to the WT line. However, in ischemia-reperfusion of the heart, the response to dobutamine was not altered between NOX5 KI and WT mice (Fig 4F). Thus, in our humanised NOX5 KI mice, neither post-ischemic outcome after permanent and/or transient heart ischemia nor long-term outcomes after hindlimb ischemia were modified. These data suggest that post-ischemic ROS formation, infarct volume and functional outcomes were all increased in a brain-specific manner in our NOX5 KI humanised model as a result of NOX5 expression.

### Human microvascular brain endothelial cells, acute post-reoxygenation leakiness is prevented by pharmacological NOX5 inhibition

Calcium is critical for tight junction function specially for characteristics of cerebral microvessels and blood-brain barrier stability. In fact, calcium overload post-stroke has been described as a key mechanism of BBB disruption by altering junction proteins [24]. Thus, we hypothesised that NOX5 is the missing mechanistic link in early BBB-dependent reperfusion injury. Moreover, thrombectomy procedures have been recently associated to reperfusion damage and early BBB disruption what may lead to hemorrhagic transformation and poor clinical outcomes. Therefore, there is a novel trend of designing experimental clinical trials to pharmacologically block BBB opening in patients undergoing reperfusion therapy of stroke [4].

To further evaluate whether NOX inhibition would be a promising candidate for this novel strategy, we used human brain microvascular endothelial cell (HBMECs) as an *in vitro* ischemia model (Fig 5A). After confluence, we subjected HBMECs to 6h of hypoxia followed by 24h of re-oxygenation and assessment of cell permeability (Fig 5B) in the presence or absence of the relatively NOX5-specific inhibitor, MI090 (0.01 μM). Since current innovative clinical approaches suggest to perform the thrombectomy procedure in presence of the pharmacological agent, we added the NOX5 inhibitor in the acute state before re-oxygenation. However, NOX4 is induced 4h post-re-oxygenation (Fig S2) although it also affects BBB disruption [14]. Then, HBMECs were treated both acutely and post-re-oxygenation with the relatively NOX4-specific inhibitor, M13 (0.2 μM) in order to mimic pre- and post-thrombectomy neuroprotective treatment. Whilst the NOX4 inhibitor was protective when added either early or late, the NOX5 inhibitor was only protective when added early (Fig 5C). These data confirm the strictly acute role of NOX5 in blood-brain-barrier breakdown post-re-oxygenation/reperfusion in stroke while NOX4 seem to play a key role in later stages. At the same time the efficacy of MI090 suggests a preventive therapeutic option, ideally as a pre-treatment or co-administration with the thrombolytic drug, i.e. rt-PA, or thrombectomy procedure.

**Fig 5.**
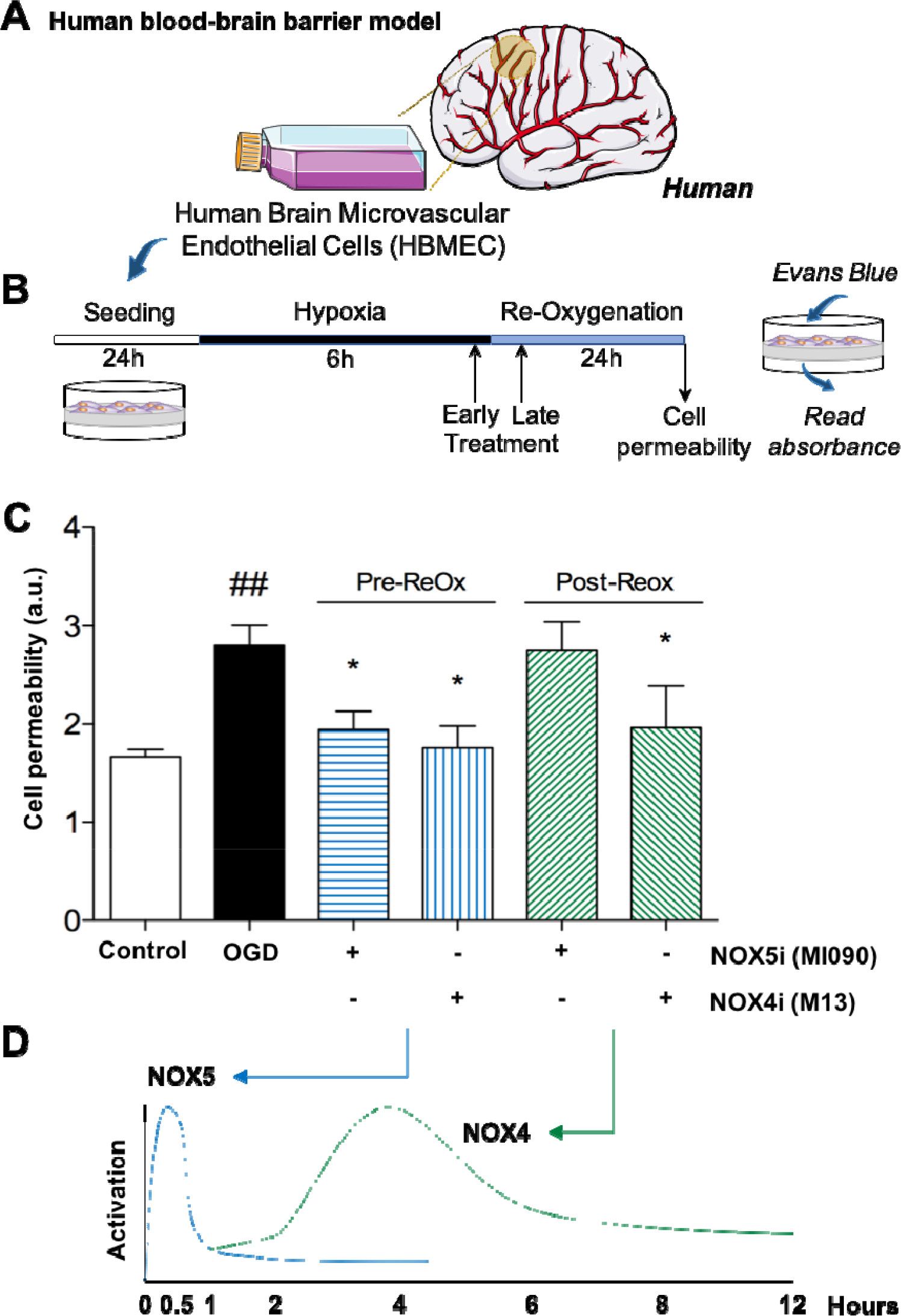
NOX5 inhibition before re-oxygenation reduced cell permeability to basal levels using a human *in vitro* ischemia model. (A) Human brain microvascular endothelial cells (HBMEC) were incubated during 24h and later seeded on trans-well inserts where cell permeability was assessed using the Evans blue dye. (B) After incubation and seeding at physiological conditions, HBMEC were subjected to 6h of hypoxia period and 24h of Re-Ox. Cells were treated with both NOX4 (M13) and NOX5 inhibitors (MI090) at early (20 min before Re-Ox) and late (20 post-Re-Ox) time-points. (C) Cell permeability was assessed by measuring Evans Blue fluorescence post-hypoxia. Evans Blue diffusion was significantly reduced in cells subjected to early treatment of MI090 (0.01µM), which preferably inhibits NOX5, and M13 (0.2 µM), mainly a NOX4 inhibitor (* P < 0.05). Contrary, late treatment was only effective after M13 (0.2 µM) treatment compared to non-treated cells (^##^ P < 0.01). (D) NOX5 activation takes place within the first 30 min/1h post-hypoxia (see figure 3) while NOX4 activation peak shows around 5h post-ischemia [15].

## Discussion

Here, we identify NOX5-derived ROS as a missing link in acute stroke therapy between the post-reperfusion calcium overload and frequent side effect of blood brain barrier breakdown and worsened outcome. NADPH oxidases in general are promising therapeutic targets in ROS-associated disease states [5,24,25]. Since most pre-clinical validation experiments are conducted in mice and rats, which lack the NOX5 gene, NOX5 has been under-studied. We provide now the first humanized NOX5 KI mouse model mimicking to a large extent the human physiological NOX5 expression pattern, i.e. endothelial and white blood cells.

In principle, our observed *in vivo* phenotype in NOX5KI mice could derive from both cell types, endothelial and white blood cells, and we can certainly not completely rule out a contribution of the latter. However, several lines of evidence support a fully sufficient role of endothelial NOX5. First, our *in vitro* mouse organotypic culture, essentially free of white blood cell contributions, and second, our HBMEC model, per definition entirely free of white blood cells, are in full agreement with a crucial endothelial location of NOX5. Third, the detrimental role of NOX5 in post-reperfusion was highly specific for brain; NOX5 played no acute role in myocardial infarction, transient heart ischemia, or in hindlimb ischemia, suggesting indirectly the BBB and its cellular components as the main location of NOX5. Moreover, endothelial NOX5 would be already at the site of action with respect to BBB breakdown, whereas white blood cells would need to adhere and only a fraction of them would be pre-exposed to post-reperfusion calcium surges.

Therapeutically, one additional aspect is worth considering, i.e. the previous identification of neuronal (and endothelial) NOX4 in stroke [14,26]. This enzyme does not contribute to the here investigated early stages of BBB opening. Therefore, any prospective therapy, however, will want to cover both isoforms, acutely induced NOX5 and sub-acutely (i.e. within 4-6 hours) induced NOX4 to further inhibit the detrimental role of NOX4 and NOX5 at different pathomechanisms stages. Thus, a combined NOX4/5 inhibitor or compound combination would be ideal.

Importantly, our data suggest that thrombolysis and thrombectomy should be conducted in the presence of NOX inhibition to minimalize the chances of a post-reperfusion, calcium- and ROS-dependent acute and deleterious opening of the blood-brain barrier. Our findings provide a clear rationale for further development of a pharmacological NOX inhibitor (ideally with combined NOX4/5 profile) as a first-in-class neuroprotective strategy after stroke, to be co-administrated already with the onset of reperfusion.

## Materials and Methods

### Animals

All animal experiments were performed according to the EU Directive 2010/63/EU for animal experiments and approved by the German Animal Welfare Act (German Ministry of Agriculture, Health and Economic Cooperation), the Dutch law on animal experiments and the institutional Ethics Committee of Universidad Autónoma de Madrid, Madrid, Spain. Animals were socially housed under controlled conditions (22°C, 55–65% humidity, 12h light-dark cycle), and could free access water and standard laboratory chow. Adult male and female mice of 8-20 weeks aged were used. NOX5 KI animals were compared to their respective matched WT line. The drop-out rates in all ischemia models (heart ischemia, hindlimb ischemia and stroke) are included in Table S1. Post-hoc power analysis for post-stroke infarct size is included in Table S2.

### Generation of the NOX5 knock-in (KI) mouse

Since the rodent genome naturally lacks the *NADPH oxidase 5* gene, we created a new mouse line expressing the human *NOX5* gene under the control of the *Tie2* promoter. The model was developed using the *hypoxanthine phospho-ribosyl-transferase* (*Hprt*) targeted transgenic approach. This locus protects transgenic constructs inserted into this region against gene silencing, positional or methylation effects, and tissue-specific promoters inserted in the *Hprt* locus conserve their expression patterns. Importantly, *Hprt*-deficient mice are phenotypically normal. Targeted insertion at the *Hprt* locus overcomes the unpredictable position effects inherent in transgenic methods relying on random integration. Thus, a respective KI line allows experiments to be conducted with a single mouse line, in contrast to the 3-5 transgenic lines usually required using classical technology.

The *Hprt* gene is localised on the X chromosome, necessitating analysis of hemizygous males or homozygous females. The targeting vector was obtained by subcloning of a transgenic targeting cassette containing the human *NOX5* beta gene under control of the *Tie2* promoter into the *Hprt* targeting vector (GenOway), upstream of the human Growth Hormone polyA sequence.

E14 ES cells of 129SV mice were transfected with the linearised targeting vector, selected with HAT (hypoxanthine-aminopterin-thymidine medium) medium and resistant clones screened (Southern blot) for presence of the 3’ and 5’ homologous recombination events. Selected ES cells were injected into C57Bl6/J blastocysts and re-implanted into OF1 (oestrus phase) pseudo-pregnant females, which resulted in chimeric males. Chimeras were mated with WT C57Bl/6 females and agouti female offspring was genotyped to confirm the presence of the recombined X chromosome. The heterozygous females were then mated with transmitting male chimeras to generate hemizygous males and homozygous females. All procedures were carried out under SPF conditions by GenOway, which houses all mice at Charles River.

### Validation of the NOX5 knock-in (KI) mouse

#### Isolation of bone marrow macrophages

Bone marrow was isolated and cells were cultured for one week in RPMI-1640 (Gibco with Glutamac, 2g/L glucose) supplemented with 10% fetal cow serum (FCS), 100U/mL Penicillin-Streptomycin, and 15% L929-conditioned medium to generate bone-marrow-derived macrophages. Cells were then dissociated with lidocaine (200mg, 1mL 0.5M EDTA in 50mL PBS) before differentiating them with EtOh, IFN-γ and IL-4 to become M0, M1 and M2a subtype macrophages respectively.

#### FACS of spleen cells

Spleen was gently dissociated through a 70 μm cell strainer (Greiner), treated with erylysis buffer, blocked with FcR-blocking (eBioscience) and stained for total leukocytes (CD45^+^, CD45 PerCP, Biolegend, Germany), total T cells (CD3^+^, CD3 eF450, eBioscience, Germany), B cells (CD19^+^, CD19 FITC, BD Bioscience, Germany), monocytes (CD11b^high^ Ly6G^-^ CD11c^-/low^, CD11b BV510 Biolegend, Ly6G APC-Cy7 BD Bioscience, CD11c PE-Cy7, eBioscience) and granulocytes (CD11b^high^ Ly6G^high^). Labeled cells were sorted on BD FACS ARIA I to >95% purity.

#### Microvascular brain endothelial cell isolation

Brain capillary endothelial cells (MBCEC) from eNOX4 KO and WT mice were isolated as described in [27,28]. After sacrificing the mice, forebrains were collected, meninges removed and the tissue was minced and digested with a mixture of 0.75mg/ml collagenase CLS2 (Worthington) and 10U/mL DNAse (Sigma-Aldrich, Germany) in Dulbecco’s modified Eagle medium (DMEM; Sigma-Aldrich, Germany) for 1h at 37°C. To remove myelin, the pellet was re-suspended in BSA-DMEM (20% w/v) and centrifuged (1000x g, 20 min). The pellet was re-suspended and further digested with 1mg/ml collagenase-dispase (Roche, Germany) and 10U/ml SNAse in DMED for 1h at 37°C. Microvascular endothelial capillaries were separated on a 33% continuous Percoll gradient, collected and plated in petri-dishes coated with collagen IV/fibronectin (Sigma-Aldrich, Germany). Cultures were maintained in DMEM supplemented with 20% plasma-derived bovine serum (First Link, Germany), 50μg/ml gentamicin (Sigma-Aldrich, Germany), 2mM L-glutamine (Sigma-Aldrich, Germany), 4μg/ml puromycin (Alexix GmbH, Grünberg, Germany) and 1ng/ml basic fibroblast growth factor (Roche, Germany).

### Organotypic hippocampal culture (OHCs) of NOX5 KI mice

Hippocampal brain slices for cultures were obtained from brains of 7- to 10-day-old WT and NOX5 KI mice. Organotypic cultures were prepared based on the methods previously described in [29,30]. Briefly, pups were quickly decapitated and brains removed from the skull and dissected. The hippocampus was cut into 300μm-thick slices using a Tissue Chopper Mcllwain. Then, they were separated in sterile ice-cold Hank’s balanced salt solution (HBSS, Biowest, Madrid, Spain) containing (in mM): glucose 15, CaCl_2_ 1.3, KCl 5.36, NaCl 137.93, KH_2_PO_4_ 0.44, Na_2_HPO_4_ 0.34, MgCl_2_ 0.49, MgSO_4_ 0.44, NaHCO_3_ 4.1, HEPES 25, 100 U/ml penicillin, and 0.100 mg/ml gentamicin. Six slices were placed on each Millicell-0.4 μm culture inserts (Millipore, Madrid, Spain) within each well of a six-well culture plate. Specific neurobasal medium (Invitrogen, Madrid, Spain) enriched with 10% of fetal bovine serum (Sigma-Aldrich, Madrid, Spain) was used for the next 24h (1 ml/well). 24h later, B27 supplement and antioxidants were added to the culture medium. Slices were in culture for 4 days before inducing the oxygen and glucose deprivation (OGD) period. On day 6, inserts were placed in 1 ml of OGD solution composed of (in mM): NaCl 137.93, KCl 5.36, CaCl_2_ 2, MgSO_4_ 1.19, NaHCO_3_ 26, KH_2_PO_4_ 1.18, and 2-deoxyglucose 11 (Sigma-Aldrich, Madrid, Spain). The cultures were then placed in an airtight chamber (Billups and Rothenberg) and exposed during 3 min to 95% N_2_/5% CO_2_ gas flow to ensure oxygen removal. After that, the chamber was sealed for 15 min at 37°C (OGD period). After 15 minutes, the cultures were returned to normal oxygen and glucose concentrations for 24h (Re-Ox period).

### *In vitro* ROS formation in organotypic hippocampal cultures

ROS production was evaluated in real-time by the fluorescence dye dihidroethidium [31,32] (Thermo Fisher Scientific, The Netherlands). A stock solution of DHE (3.2 mM) was dissolved in Krebs solution and added to the culture insert. Fluorescence measurements were performed at 0, 15 min, 30 min, 1h, 2h and 4h after the OGD period in both WT and NOX5 KI mice using a 10X objective in the CA1 region of the hippocampus. Same emission and excitation wavelength were used. Fluorescence analysis was performed using the Metamorph software version 7.0.

### Ca^2+^ overload in organotypic hippocampal cultures

Hippocampal brain slices were in culture 4 days as previously described. To induce Ca^2+^ overload, 10 µM of ionophore A23187 (Sigma-Aldrich, The Netherlands) was directly added to the culture medium and ROS formation was subsequently measured objective in the CA1 region of the hippocampus at 0, 15 and 30 min post-A23187 addition. Fluorescence analysis was performed using the Metamorph software version 7.0. Different time-points were selected based on previously described ROS kinetics.

### Transient occlusion of the middle cerebral artery (tMCAO model)

The model has previously been established as described in [26]. Animals were anesthetized with isoflurane (1,5-2% in oxygen). The animal was placed on a heating-pad, and rectal temperature was maintained at 37.0°C using a feedback-controlled infrared lamp. Focal cerebral ischemia was induced using an intraluminal filament technique. Using a surgical microscope (Wild M5A, Wild Heerbrugg, Gais, CH), a midline neck incision was made and the right common and external carotid arteries were isolated and permanently ligated. A microvascular clip was temporarily placed on the internal carotid artery. A silicon rubber-coated 6.0 nylon monofilament (602312PK10, Doccol Corporation, Sharon, MA, USA) for mice was inserted through a small incision into the common carotid artery and advanced into the internal carotid artery until a resistance is felt. The tip of the monofilament should be located intracranially at the origin of the right middle cerebral artery and thereby interrupting blood flow. The filament was held in place by a tourniquet suture that has been prepared before to prevent dislocation during the ischemia period and the wound was closed. Reperfusion was initiated 30 minutes after occlusion. After the surgery, wounds were carefully sutured and animals recovered from surgery in a temperature-controlled environment. Animals were excluded from the stroke analysis if animals died before the predefined experimental end-point, if an intracerebral hemorrhage occurred or if the animal scored 0 on the Bederson score.

### Brain infarct volume measurements

The ischemic lesion was measured 24 hours after tMCAO using TTC staining [33]. The brain was cut in three 2 mm thick coronal sections using a mouse brain slice matrix (Harvard Apparatus, Holliston, MA, USA). The slices were soaked for 10 min in a freshly-prepared solution of 2% 2,3,5-triphenyltetrazolium hydrochloride (TTC, Sigma-Aldrich, Germany/The Netherlands). Total indirect (i.e corrected for brain edema) infarct volume was calculated by volumetry (ImageJ 1.49 software, National Institutes of Health) according to the formula: V_indirect_ (mm^3^) = V_infarct_ x (1-(V_ih_-V_ch_) / V_ch_, where the term V_ih_-V_ch_ represents the volume difference between the ipsilateral and contralateral hemisphere and (V_i_-V_c_) / V_c_ expresses this difference as % of the control hemisphere.

### Neurological behaviour

The mice were assessed for neurological behaviour just before sacrifice to determine the final functional status. Neurological deficits were measured in a blinded manner on a 0 to 5 scale using the Bederson Score [33] with the following definitions: Score 0, no apparent neurological deficits; 1, body torsion and forelimb flexion; 2, right side weakness and thus decreased resistance to lateral push; 3, unidirectional circling behaviour; 4, longitudinal spinning; 5, no movement.

### Motor function

Prior to sacrifice, the mice were also scored for neurological motor deficits according to the Grip Test [26]. Each mouse was given a discrete value from 0 to 5. This score is used to evaluate motor function and coordination. The apparatus is a metal rod (0.22 cm diameter, 50cm length) between two vertical supports at a height of 40 cm over a flat surface. The animal is placed mid-way on this rod and is rated according to the following system: Score 0, falls off; 1, hangs on to string by one or both fore paws; 2, as for 1, and attempts to climb on to string; 3, hangs on to string by one or both fore paws plus one or both hind paws; 4, hangs on to string by fore and hind paws plus tail wrapped around string; 5, escape (towards the supports).

### Blood-brain barrier function

To determine the permeability of the cerebral vasculature and brain edema, 2% Evans blue tracer (Sigma-Aldrich, Germany/The Netherlands) diluted in 0.9% NaCl was injected i.p. at reperfusion. Measurement of Evans Blue extravasation was performed as described in [26].

### Oxidative stress: DHE staining

The presence of ROS was determined using dihydroethidium (Sigma-Aldrich, Germany, stock solution 2mM) staining in coronal brain sections taken from identical regions (-0.5mm from bregma) of the different animal groups. Briefly, frozen sections were incubated in 2μM DHE for 30 minutes at 37°C, washed three times with PBS and incubated with Hoechst (Hoechst 33342, Sigma-Aldrich, Germany) 2 ng/ml for 10 min. All sections were analyzed and acquired with a Nikon Eclipse 50i microscope equipped with the DS-U3 DS camera control unit. The relative pixel intensity was measured in identical regions with NIS-Elements software (Nikon, Tokyo, Japan). Digital images were processed using Adobe Photoshop (Adobe Systems, San Jose, CA).

### Heart ischemia: Myocardial infarction and ischemia reperfusion

Mice aged 8-16 weeks were subjected to permanent or transient ligation of the left descending coronary artery. After administration of an analgetic (buprenorphine s.c. 0,05mg/kg, Temgesic, Schering-Plough), mice were anaesthetized with isoflurane (Abbott forene Isoflurane) 4-5% in air and intubated per orally with a stainless-steel tube. The tube was connected to a respirator (rodent ventilator Microvent type 845, Hugo Sachs Electronic, Germany), set at a stroke volume of 250μL and a rate of 210 strokes/min. Anaesthesia was then maintained with 2-3% isoflurane in air via a vaporizor (Univentor, UNO Roestvaststaal BV) connected to the respirator. The mouse was placed on a heating pad (UNO temperature control unit, UNO Roestvaststaal BV) and body temperature was monitored using a rectal probe and maintained at 37.0°C using a feedback-controlled infrared light. During surgery, an ECG was recorded with IDEEQ software (IDEE, Maastricht University). A left thoracotomy was performed to expose the heart. Then, the left descending coronary artery (LAD) was ligated with a 6-0 polypropylene suture (Surgipo, Chicago, IL, USA), just proximal to its main branching point. The suture was tied around a 3 mm-long polyethylene tube (PE-10) to induce ischemia. Ischemia was assessed by a discolouration of the tissue (from red to pink-white) and ST-elevation on ECG. After 45 minutes, the blood flow was re-established by removal of the polyethylene-tube. The occurrence of reperfusion was assessed by the colour of the tissue turning red again. For the myocardial infarct model, the LAD was ligated with a 6-0 polypropylene suture permanently. The chest was closed with 5-0 silk sutures (Ethicon). The animals were weaned from the respirator and the endotracheal tube was removed, once the mice breathed spontaneously. After surgery, mice were allowed to recover at thermoneutral temperature (28°C). In the morning and afternoon of the day after, s.c. 0.05 mg/kg buprenorphine hydrochloride was repeated to relieve pain.

### Evaluation of infarct size in heart

After the hemodynamic measurements at day 28, the heart was excised and the atria were removed. The ventricles were cut transversally at 3 mm from the apex. The apical part was fixed in formalin and processed for paraffin embedding. Paraffin sections of 4 μm were stained with AZAN (infarct size) or Picrosirius red (collagen content). Images were visualised under light microscopy (Zeiss Axioskop microscope) and digitalised using a Leica DFC490 camera (Leica Microsystems Ltd, Heerbrugg) Pictures were analysed using the Leica Qwin pro v3.5.1 software. Infarct sizes are expressed as percentage area of the total left ventricular tissue area. Animals with no visible infarct were deleted from all analysis.

### Ligation of the femoral artery

Directly after a basal Doppler measurement, anaesthesia was maintained with 1,5-2% isoflurane. The mouse was placed on the back on a heating pad (UNO temperature control unit, UNO Roestvaststaal BV) and body temperature was monitored using a rectal probe and maintained at 37.0°C using a feedback-controlled infrared light. The right groin was disinfected and the right femoral artery was ligated by placing a suture (5-0 silk) around the femoral artery in between the branching of the a. epigastrica and the a. poplitea. These last two arteries were also ligated to prevent collateral flow and backflow respectively. The wound was then closed with a 4-0 polysorb suture and the mouse was allowed to recover. Discoloration of the paw was visible after the ligating surgery and an accompanying drop in blood flow was visible with laser Doppler.

### Capillary density

After the last Doppler measurement at day 28, the musculus adductor and musculus gastrocnemius were prepared free, dissected and formalin fixed. Paraffin embedded sections of 4 μm were stained for capillary cells, using a monoclonal rat anti-mouse antibody to CD31 (PECAM-1) (Histonova-Dianova, Cat. no DIA310) diluted 1:50. As secondary antibody, biotin labelled rabbit anti-rat antibody (dakocytomotion Denmark no. E0468) was used diluted 1:200. Images were visualised under light microscopy (Zeiss Axioskop microscope) and digitalised using a Leica DFC490 camera (Leica Microsystems Ltd, Heerbrugg). Pictures were analysed using the Leica Qwin pro v3.5.1 software. For each animal, three random pictures were taken per muscle sample and the amount of capillaries is expressed as number per mm^2^.

### Human brain microvascular endothelial cells (HBMECs) subjected to hypoxia

HBMEC (Cell systems, USA) between passage 3 and 9 were cultured to approximately 95% confluence using recommended cell medium (EGM-2 MV BulletKit, Lonza, The Netherlands) enriched with 5% fetal bovine serum (FBS; Sigma-Aldrich, The Netherlands) before starting the experiment. For hypoxia studies, HBMECs were always seeded at specific cell density (6×10^4^ cells/ml) in 12 wells-plate and incubated during 24h at 37°C. Then, cell medium was replaced with non-FBS enriched medium (2 ml/well) following by 6h of hypoxia (94,8% N_2_, 0.2% O_2_ and 5% CO_2_) at 37°C using hypoxia workstations (Ruskin Invivo2 400 station, The Netherlands). The hypoxia period was followed by 24h of re-oxygenation at 37°C, enriched medium and normoxia conditions (75% N_2_, 20% O_2_ and 5% CO_2_) in the presence or absence of treatment. All culture surfaces were pre-treated with fibronectin solution (1:100 in PBS, Sigma-Aldrich, The Netherlands).

HBMECs were treated at two different time points: early treatment (25 min before re-oxygenation) and late treatment (25 min after re-oxygenation) with 0,01μM MI090 (NOX5 inhibitor) and 0,2μM M13 (NOX4 inhibitor, Glucox, Sweden).

### Assessment of cell permeability in HBMECs

2 x 10^4^ HBMECs were seeded and incubated during 24h on Transwell inserts (collagen-coated Transwell Pore Polyester Membrane Insert; pore size = 3.0 µm, Corning, The Netherlands) before inducing 6h of hypoxia (94,8% N_2_, 0.2% O_2_ and 5% CO_2_) followed by 24h re-oxygenation period in presence/absence of early or late treatment. The Evans Blue dye (Sigma-Aldrich, The Netherlands) was used to assess cell permeability on the insert. Before the diffusion experiment, the medium was removed and cells were washed once with assay buffer (37°C-warm PBS). 1.5 ml of the same buffer was added to the ab-luminal side of the insert. Permeability buffer (0.5 ml) containing 4% bovine serum albumin (Sigma-Aldrich, The Netherlands) and 0.67 mg/ml Evans blue dye in PBS was loaded on the luminal side of the insert followed by 15 min incubation at 37°C. Evans Blue concentration in the ab-luminal chamber was measured by determining the absorbance of 150 µl buffer at 630 nm using a spectrophotometer.

### Statistics

All data are expressed as mean ± SEM. Using the GraphPad Prism 6.0 software package data were assessed for normal distribution using the D’Agostino & Person omnibus normality test. For each outcome parameter, outliers clearly >2SD were not considered in the analysis. Statistical differences between mean continuous values were determined by Student’s two-tailed t-test. For repeated measurements, a two-way ANOVA was used. Categorical values or continuous values that did not pass the normality test were assessed using the Mann-Whitney t-test. A value of p<0.05 was considered as statistically significant.

## Acknowledgements

This study was supported by an ERC Advanced Investigator Grant (to HHHWS), Spanish Ministry of Economy and Competence Ref. SAF2015-63935R (to MGL), the Deutsche Forschungsgemeisnchaft (C.K.), Fondo de Investigaciones Sanitarias (FIS) (ISCIII/FEDER) (Programa Miguel Servet: CP14/00008, PI16/00735) and Fundación Mutua Madrileña (J.E.), Spanish Ministry of Economy and Competence Ref. SAF2015-63935R (M.G.L.) and short-term scientific missions by the COST Action EU-ROS (to AC). A. Brouns-Strzelecka and J. Debets are gratefully acknowledged for performing the heart ischemia *in vivo* experiments. H. van Essen is gratefully acknowledged for performing the blood pressure telemetry experiments.

## Author contributions

**Conceptualization**: Harald Schmidt

**Formal analysis**: Ana I Casas, Pamela Kleikers, Eva Geuss, Friederike Langhauser

**Funding acquisition**: Harald Schmidt, Christoph Kleinschnitz, Manuela G Lopez

**Investigation**: Ana I Casas, Pamela Kleikers, Eva Geuss

**Methodology**: Ana I Casas, Pamela Kleikers, Eva Geuss, Javier Egea

**Supervision**: Harald Schmidt, Christoph Kleinschnitz, Manuela G Lopez

**Writing**: Ana I Casas, Pamela Kleikers, Harald Schmidt, Manuela G Lopez

## Competing financial interests

The authors declare no competing financial interests.

## Supporting Information - PLOS Biology

### Supplementary Figures

**Fig S1.**
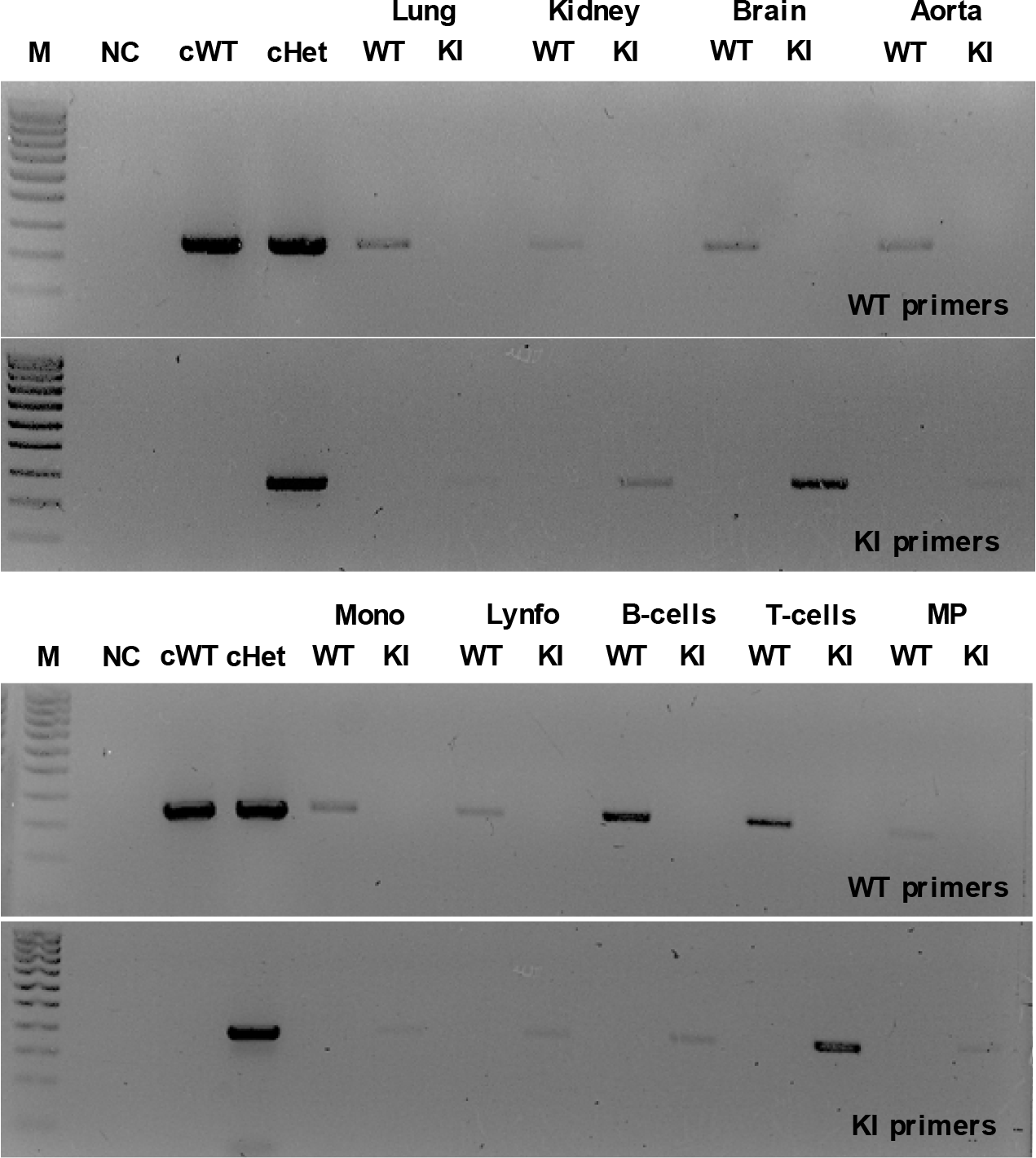
NOX5 KI mice genotyping. Control (WT) and NOX5 KI mice tail genomic DNA were purified and a PCR was performed to amplify the *Nox5* sequence in different tissues such as lung, kidney, brain, aorta and white blood cells (monocytes, lymphocytes, B- and T-cells). Nox5 DNA sequence were detected in NOX5 KI mice while no signal was shown in WT mice.

**Fig S2.**
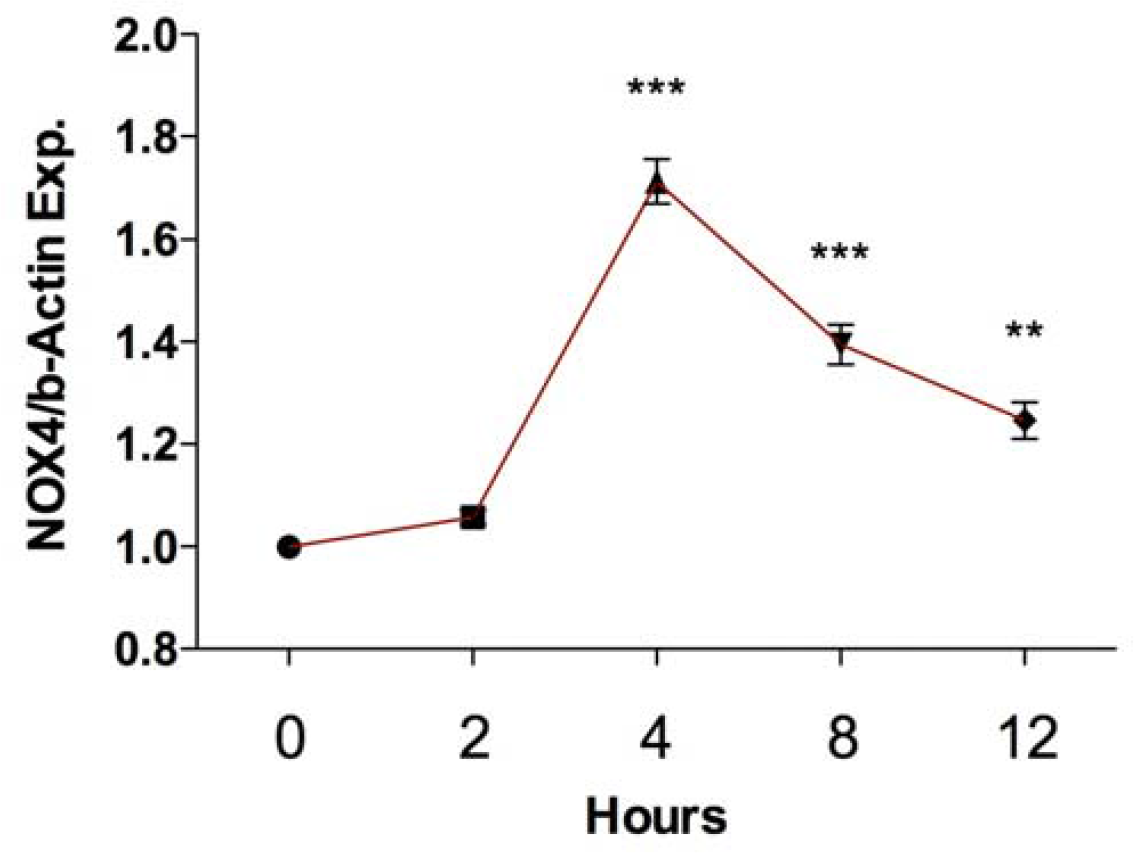
NADPH oxidases 4 (NOX4) are up-regulated at different time-points after oxygen and glucose deprivation (OGD). Organotypic hippocampal cultures (OHCs) prepared from mice hippocampal brain slices were cultured for 4 days and subsequently subjected to 15 min of OGD period. Brain slices were collected at 0, 2, 4, 8, and 12 after OGD for later gene expression analysis. NOX4 expression was up-regulated at 4h, 8h and 12h in comparison with the beginning of the ischemia period (^**^*p* < 0.01, ^***^*p* < 0.001, n = 3).

### Supplementary tables

**Table 1.** Study design. Animals.

| Model | Measurement | Strain | Drop-out | Outliers | N |
| --- | --- | --- | --- | --- | --- |
| Hindlimb ischemia | *Doppler* | WT | 0 | 0 | 13 |
|  |  | KI | 0 | 3 | 10 |
|  | *Capillary density gastrocnemius* | WT | 0 | 0 | 13 |
|  |  | KI | 0 | 1 | 13 |
|  | *Capillary density adductor* | WT | 0 | 1 | 12 |
|  |  | KI | 0 | 0 | 14 |
| Heart ischemia permanent | *Infarct size* | WT | 12 | 0 | 16 |
|  |  | KI | 12 | 1 | 15 |
|  | *DFP* | WT | 12 | 1 | 13 |
|  |  | KI | 12 | 1 | 15 |
|  | *DFN* | WT | 12 | 1 | 13 |
|  |  | KI | 12 | 1 | 15 |
|  | *Ejection fraction* | WT | 12 | 0 | 16 |
|  |  | KI | 12 | 0 | 14 |
| Heart ischemia transient | *Infarct size* | WT | 1 | 1 | 13 |
|  |  | KI | 1 | 0 | 17 |
|  | *DFP* | WT | 1 | 1 | 12 |
|  |  | KI | 1 | 1 | 17 |
|  | *DFN* | WT | 1 | 1 | 12 |
|  |  | KI | 1 | 2 | 17 |
|  | *Ejection fraction* | WT | 1 | 1 | 12 |
|  |  | KI | 0 | 0 | 16 |
| Blood pressure | *All measurements* | WT | 2 | 0 | 19 |
|  |  | KI | 0 | 0 | 20 |
| Stroke | *Infarct size* | WT | 5 | 1 | 27 |
|  |  | KI | 5 | 4 | 25 |
|  | *Blood-brain barrier* | WT | 0 | 1 | 22 |
|  |  | KI | 3 | 2 | 26 |

*WT, wild type; KI, knock-in.*

**Table 2.**
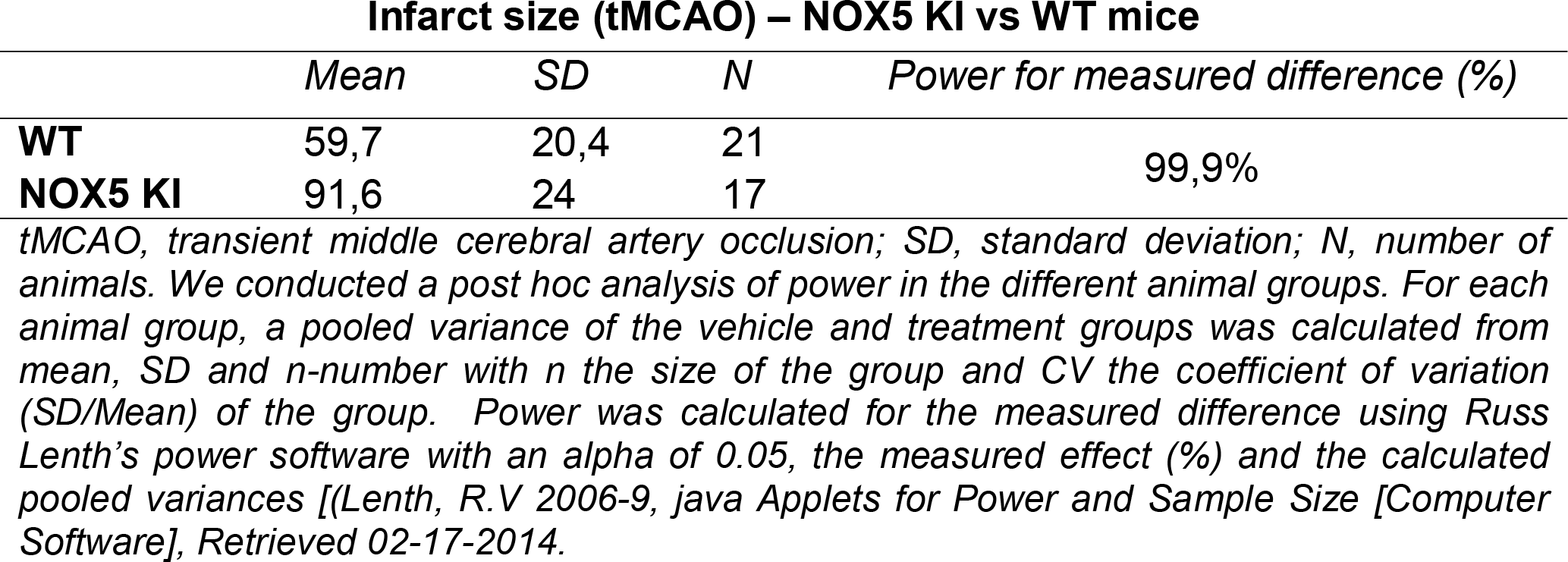
Study design. Power analysis.

